# Interface-aware molecular generative framework for protein-protein interaction modulators

**DOI:** 10.1101/2023.10.10.557742

**Authors:** Jianmin Wang, Jiashun Mao, Chunyan Li, Hongxin Xiang, Xun Wang, Shuang Wang, Zixu Wang, Yangyang Chen, Yuquan Li, Kyoung Tai No, Tao Song, Xiangxiang Zeng

## Abstract

Protein-protein interactions (PPIs) play a crucial role in numerous biochemical and biological processes. Although several structure-based molecular generative models have been developed, PPI interfaces and compounds targeting PPIs exhibit distinct physicochemical properties compared to traditional binding pockets and small-molecule drugs. As a result, generating compounds that effectively target PPIs, particularly by considering PPI complexes or interface hotspot residues, remains a significant challenge. In this work, we constructed a comprehensive dataset of PPI interfaces with active and inactive compound pairs. Based on this, we propose a novel molecular generative framework tailored to PPI interfaces, named GENiPPI. Our evaluation demonstrates that GENiPPI captures the implicit relationships between the PPI interfaces and the active molecules, and can generate novel compounds that target these interfaces. Moreover, GENiPPI can generate structurally diverse novel compounds with limited PPI interface modulators. To the best of our knowledge, this is the first exploration of a structure-based molecular generative model focused on PPI interfaces, which could facilitate the design of PPI modulators. The PPI interface-based molecular generative model enriches the existing landscape of structure-based (pocket/interface) molecular generative model.

## Introduction

A vast network of genes is interconnected through protein-protein interactions (PPIs), which are critical components of nearly every biological process under physiological conditions and are ubiquitous in various organisms and biological pathways[1–4]. Modulating PPIs broadens the drug target space and holds significant potential in drug discovery. In humans, the estimated size of the interactome ranges from 130,000 to 930,000 binary PPIs[5–7]. Despite considerable efforts, developing modulators of PPI targets, particularly those targeting PPI interfaces, remains challenging[6, 8–12]. Structure-based rational design plays a vital role in identifying lead compounds for drug discovery[13–17]. Traditional drug targets and PPI targets exhibit distinct biochemical characteristics (**Table 1**)[11, 18–22], leading to differences in the physicochemical and drug-like properties of conventional drugs and PPI modulators (**Table 1**)[11, 23–30]. Given these differences, developing molecular generative models tailored to different paradigms is crucial for designing drugs for various target types[10, 19, 31].

**Table 1.**
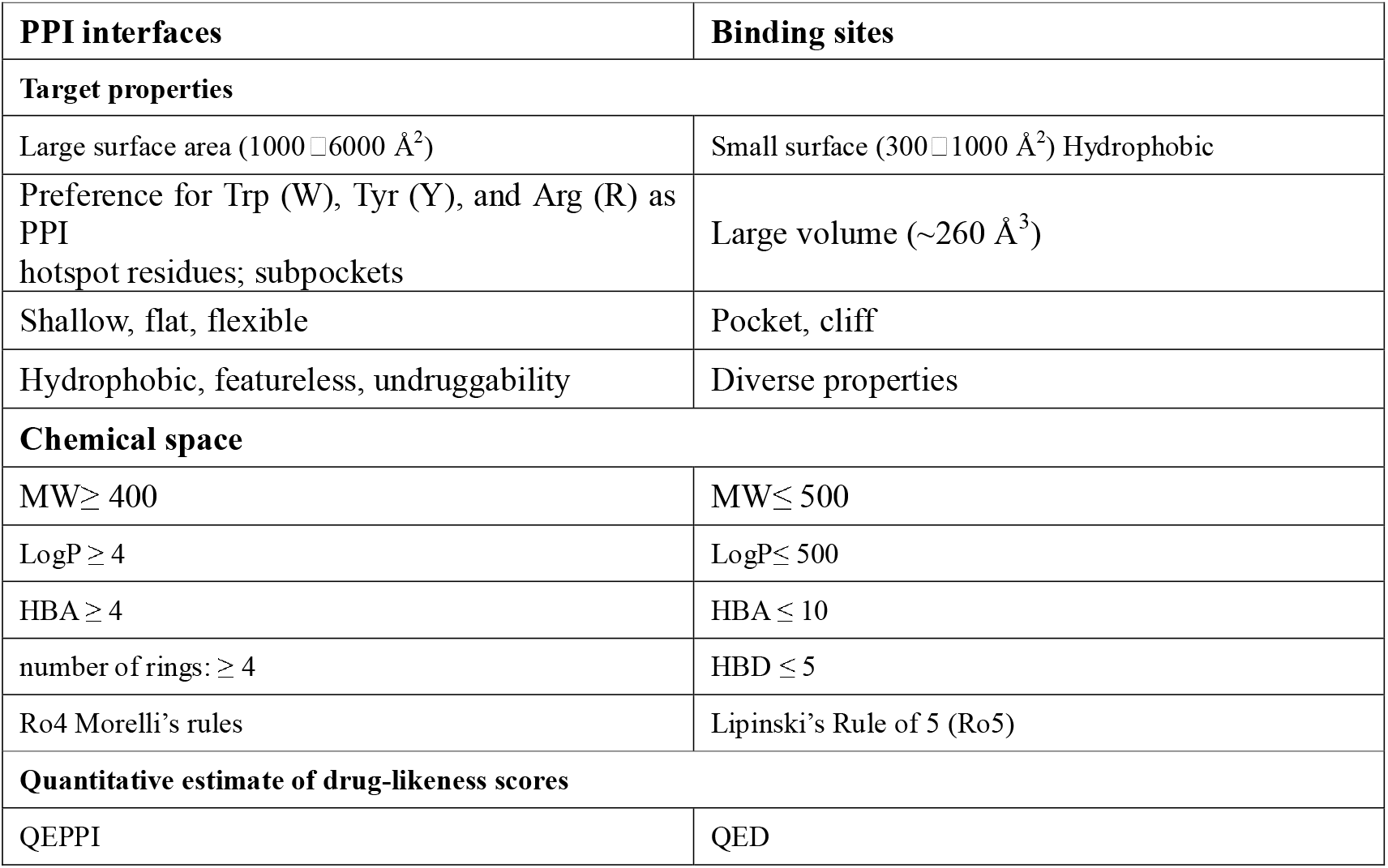
Comparisons between PPI interfaces and binding sites.

Generative artificial intelligence (AI) is capable of modeling the distribution of training samples and generating novel samples[32, 33]. In drug discovery, generative AI can accelerate the process of drug discovery by generating novel molecules with desired properties. Several excellent review articles have summarized development in this field[16, 17, 34–41]. Molecular generative models in drug design can be broadly categorized into three categories: ligand-based molecular generative (LBMG) models, structure-based molecular generative (SBMG) models (focusing on pockets or binding sites), and fragment-based molecular generative (FBMG) models. Among these, SBMG models have garnered significant attention[17, 39, 42]. While key methods for structure-based molecular generative models are well-documented[43– 51], molecular generative models specifically targeting PPI structures or interfaces are rarely reported in the literature. In recent years, classical machine learning[52–54], active learning[55], and deep learning-assisted approaches have been explored to improve the screening and design of PPI modulators[56], and some ligand-based molecular generative models for PPI modulators have been reported[57]. However, structure-based molecular generative models for PPI targets remain underexplored.

In this study, we developed GENiPPI, a structure-based conditional molecular generative framework designed for the generation of protein-protein interaction (PPI) interface modulators. The framework begins by utilizing Graph Attention Networks (GATs) to capture the subtle atomic-level interaction features present at the protein complex interface. Convolutional Neural Networks (CNNs) are then employed to extract compound representations in voxel and electron density space. Following this, a Conditional Wasserstein Generative Adversarial Network (cWGAN) integrates these features to train a model that generates compound representations targeting PPI interfaces. Finally, the LSTM network decodes the molecular embeddings into SMILES strings. The framework is designed to capture the relationship between PPI interface with active/inactive compounds, enabling the training of conditional molecular generative models specifically tailored to PPI interfaces (**Figure 1**). Conditional evaluation shows that the GENiPPI framework effectively captures the implicit relationships between PPI interfaces and active compounds, generating compounds with drug-like properties that resemble those of active compounds targeting specific PPI sites. In terms of performance, GENiPPI outperforms other generative models such as LatentGAN and ORGAN, demonstrating superiority in the novelty, diversity, and validity of the generated molecules. Additionally, in few-shot molecular generation, GENiPPI successfully generated compounds targeting the Hsp90-Cdc37 interaction with chemical properties similar to known disruptors, proving effective even with limited labeled data. In conclusion, GENiPPI represents a potent deep learning framework for structure-based design of PPI modulators.

**Figure 1.**
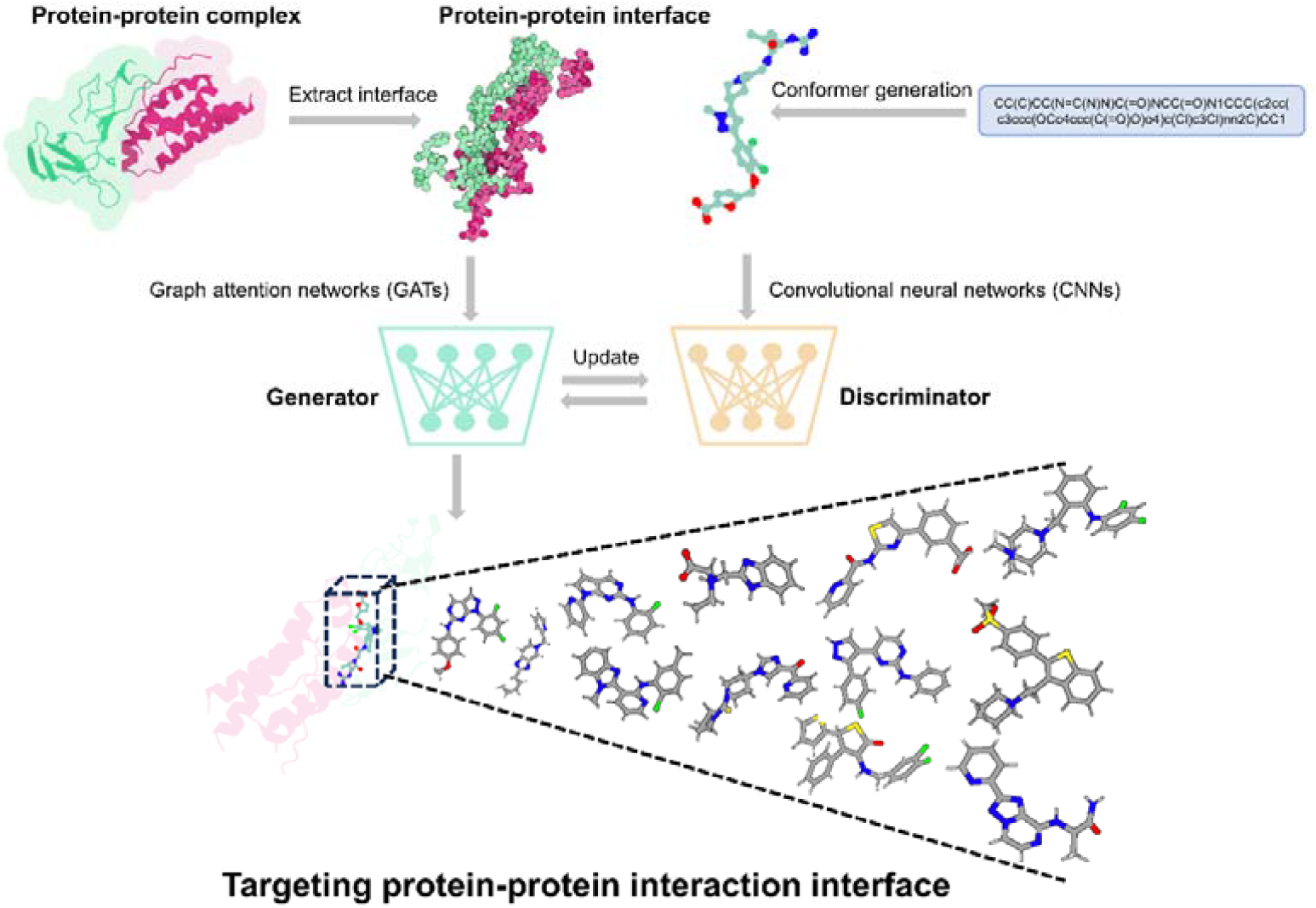
The generation of molecules targeting PPI. The 3D structural information of the protein-protein complex interface is represented as a graph, with feature representation of the interface region captured using a graph attention neural networks (GATs). The voxel and electron density of the compound are encoded by 3D convolutional neural networks (CNNs). Conditional Wasserstein generative adversarial networks (cWGAN) is trained to generate molecular embeddings conditioned on interface features. The generator takes interface features and random noise vectors to generate molecular embeddings, while the discriminator evaluates the probability that a molecule is real or generated. The condition regulates the generation of molecules constrained by specific protein-protein interfaces. Finally, long short-term memory (LSTM) networks decode the molecular representations into SMILES strings.

## Results and discussion

### Generation of molecules targeting the PPI interface

In this study, we introduce GENiPPI, a modular deep learning framework for the design of structure-based PPI modulators (**Figure 1**). GENiPPI consists of four main modules: a Graph Attention Networks (GATs) module for representation learning of the protein complex interface[58–60], a Convolutional Neural Networks (CNNs) module for molecular representation learning, Conditional Wasserstein GAN (cWGAN) module for conditional molecular generation[61], and a molecular captioning network module for decoding molecular embeddings into SMILES strings (**as shown in Supplementary Figs. 1, 2, 3, and 4, respectively**).

**Figure 2.**
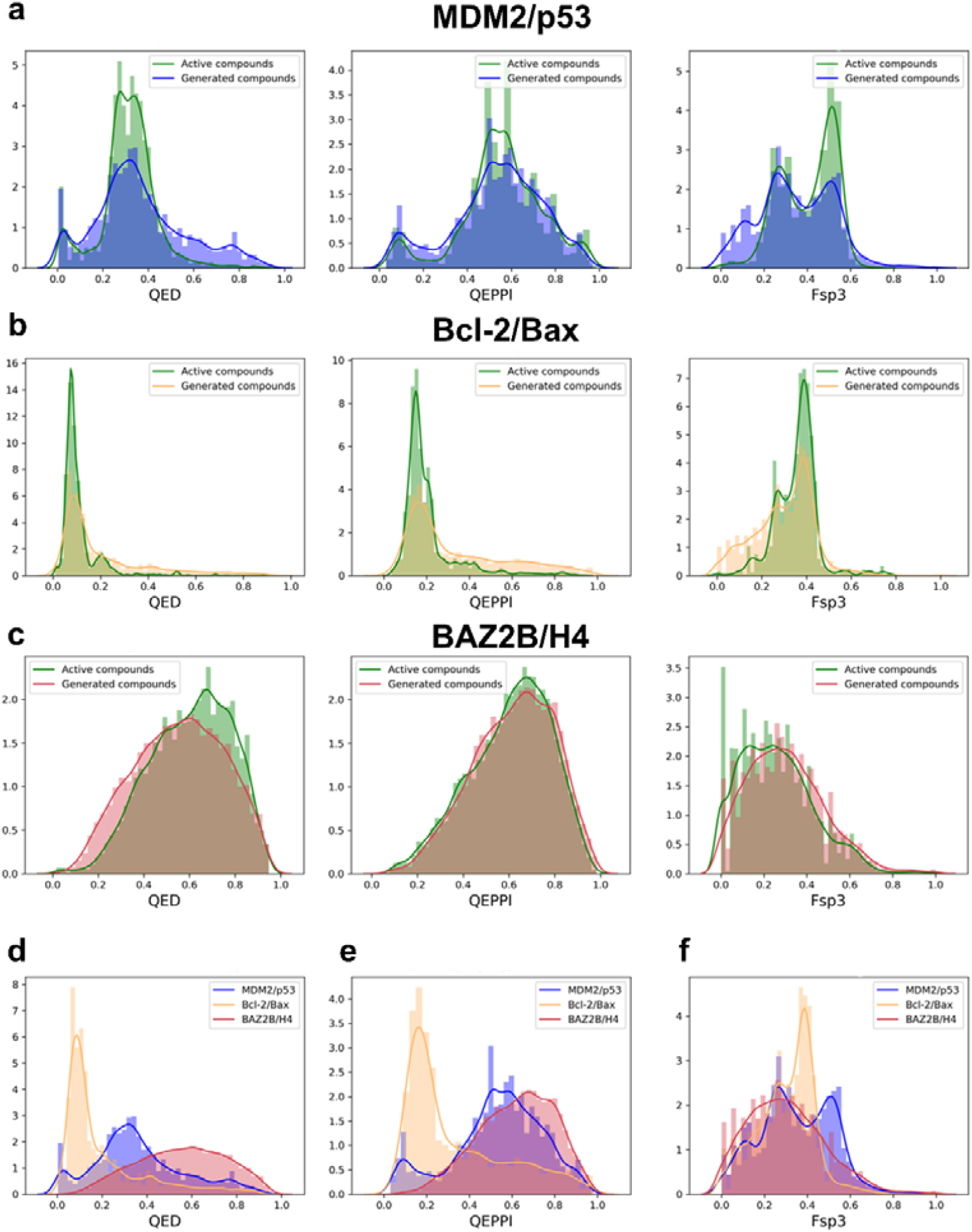
Results of conditional evaluation. (a) The distribution of QED, QEPPI and Fsp3 for active compounds and compounds generated by the GENiPPI framework for MDM2/p53; (b) The distribution of QED, QEPPI and Fsp3 for active compounds and compounds generated by the GENiPPI framework for Bcl-2/Bax; (c) The distribution of QED, QEPPI and Fsp3 for active compounds and compounds generated by the GENiPPI framework for BAZ2B/H4; (d) The QED distribution of compounds generated by the GENiPPI framework for MDM2/p53, Bcl-2/Bax and BAZ2B/H4; (e) The QEPPI distribution of compounds generated by the GENiPPI framework for MDM2/p53, Bcl-2/Bax and BAZ2B/H4; (f) The Fsp3 distribution of compounds generated by the GENiPPI framework for MDM2/p53, Bcl-2/Bax and BAZ2B/H4.

**Figure 3.**
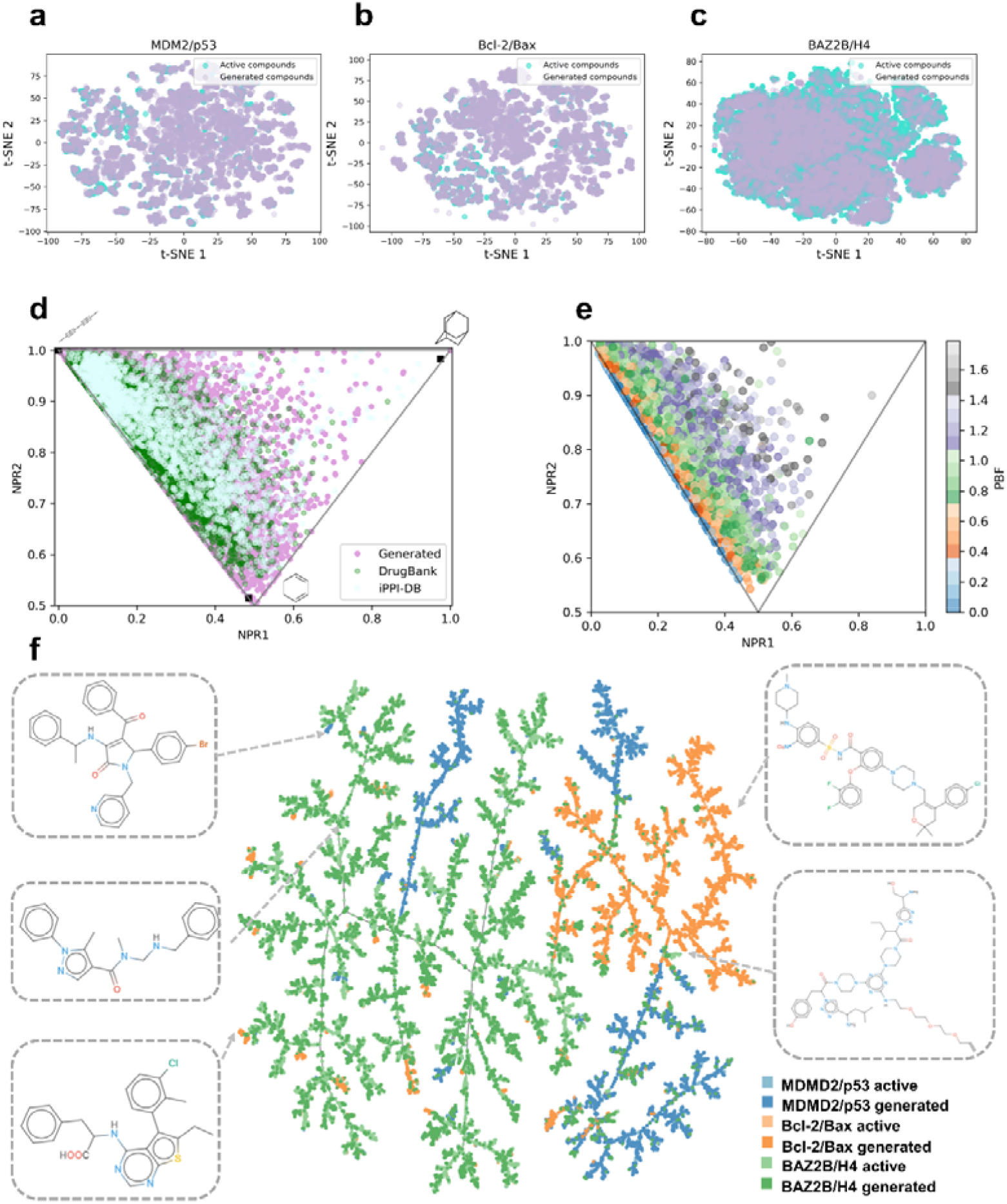
Chemical space exploration. (a) t-SNE visualization of active and generated compounds for MDM2/p53; (b) t-SNE visualization of active and generated compounds for Bcl-2/Bax; (c) t-SNE visualization of active and generated compounds for BAZ2B/H4; (d) PMI ternary density plots of generated compounds, small molecule drugs from DrugBank, and iPPI-DB inhibitors. Top left: propyne, bottom: benzene, and the top right: adamantane; (e) Molecular three-dimensionality distribution of generated molecules visualized using NPR and PBF descriptors. (f) TMAP visualization of active and generated compounds for MDM2/p53, Bcl-2/Bax and BAZ2B/H4.

**Figure 4.**
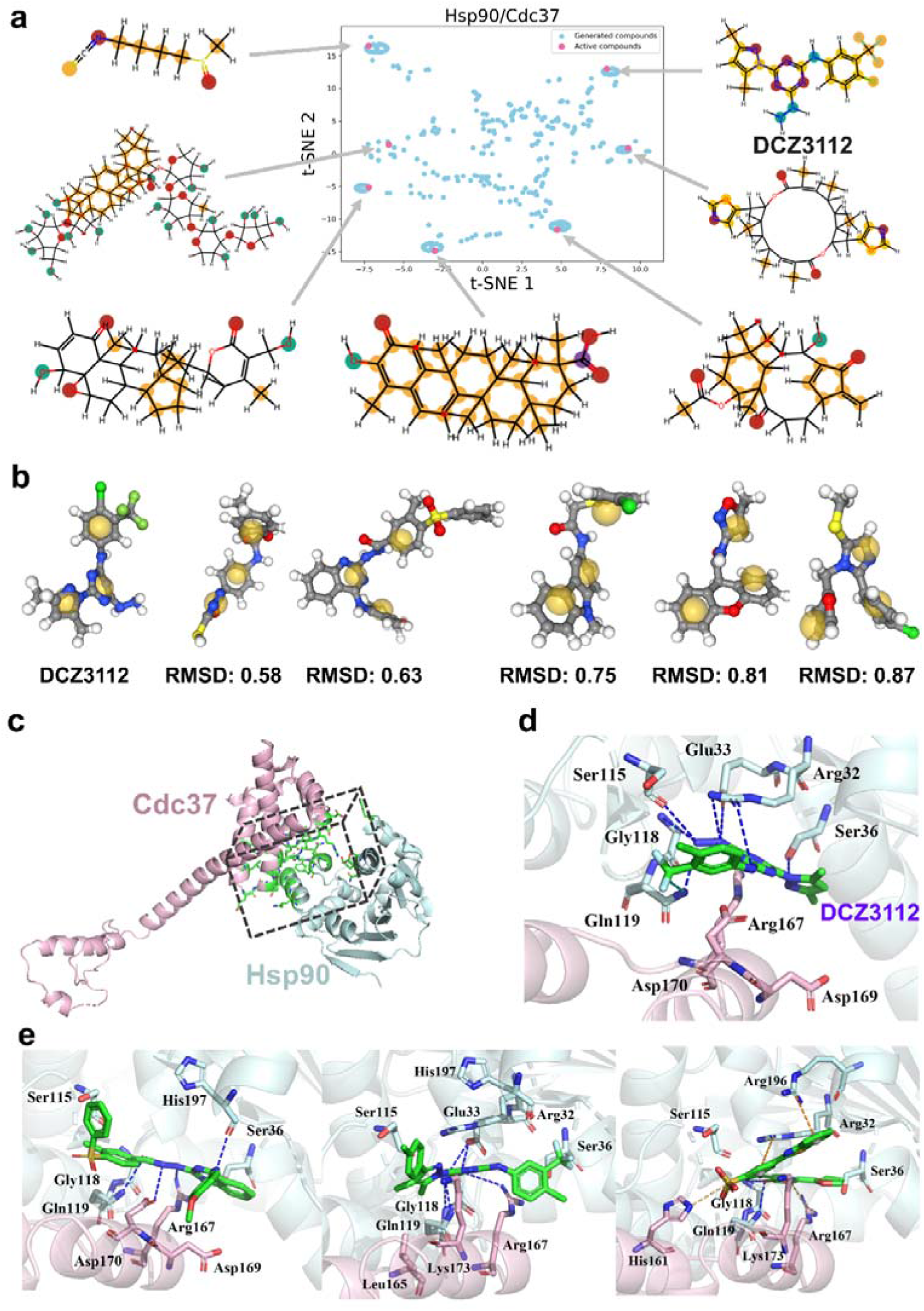
Few shot molecular generation analysis. (a) t-SNE visualization of the distribution of active and generated compounds for Hsp90/Cdc37; (b) Comparison of the pharmacophore of the generated molecules with the reference molecule(DCZ3112); (c) PPI interface region(green) of the Hsp90(palecyan)/Cdc37(lightpink) complex; (d) The complex structure of DCZ3112(green) and Hsp90(palecyan)-CDC37(lightpink) modeled by molecular docking (PDB ID: 1US7); (e) The binding poses of generated compounds(green) and Hsp90(palecyan)-CDC37(lightpink) modeled by molecular docking (PDB ID: 1US7). Hydrogen bonds are displayed as blue dotted lines, π-cation interactions are displayed as orange dotted lines.

Our framework follows four steps to generate molecules targeting PPI interfaces. First, the GATs module extracts atomic-level interaction characteristics of the protein complex interface region, effectively capturing the nuanced structural characteristics crucial for interaction. Then, the CNN module encodes the molecular features in a three-dimensional space, incorporating voxel and electronic density information [62]. This ensures a robust representation of the molecular structure that is suitable for generational tasks.

Next, the cGAN module then generates compounds targeting PPI interfaces by utilizing features from the protein complex interface to regulate the inputs[63]. This cGAN module consists of three components: the generator, the discriminator, and conditional network. The generator takes a Gaussian random noise vector and the protein complex interface features to generate a vector in the molecular embedding space, The discriminator determines whether the generated molecular embedding corresponds to a real or generated molecule, while the conditional network assesses whether the molecular embedding matches the protein complex interface features. Finally, the molecular captioning network decodes the molecular embeddings into SMILES strings. This network comprises a 3D CNN that processes the molecular embedding followed by an LSTM (Long Short-Term Memory) network, which sequentially decodes the learned embeddings into valid molecular structures in SMILES strings[64]. This step ensures that the generated molecular designs are usable in further drug design studies.

### Conditional evaluation

To comprehensively assess the validity of the conditions used as conditional molecular generative models targeting the protein complex interfaces. To do this, we conducted a detailed analysis using three distinct PPI targets: MDM2(mouse double minute 2)/p53, Bcl-2(B-cell lymphoma 2)/Bax (Bcl-2 associated X), and BAZ2B(Bromodomain adjacent to zinc finger domain protein 2B)/H4(histone). These targets were selected due to the availability of high-quality labeled data and their significance in cancer biology.

For each PPI target, we used the GENiPPI framework to generate 10,000 validated molecules and calculated the key drug-like metrics of the generated compounds: QED[27], QEPPI[28, 29] and Fsp3(fraction of sp3 carbon atoms)[65]. These metrics are essential indicators of drug-likeness, PPI-targeting drug-likeness, and molecular complexity, respectively. The aim of these calculations was to determine how well the generated molecules align with the drug-like properties of known active compounds and to evaluate the influence of conditional input on the generative process.

We then compared the QED, QEPPI, and Fsp3 distributions of the active compounds and generated compounds for MDM2/p53, Bcl-2/Bax and BAZ2B/H4 **(Figure 2)**. The results show that the drug-like property distributions of the generated compounds closely resemble those of the active compounds for all three PPI interface targets (**Figure 2a, 2b, and 2c**). This suggests that the conditional input derived from the PPI interface features plays a critical role in guiding the generation process toward biologically relevant compounds.

Interestingly, we observed differences in drug-like property distributions between the generated compounds across different PPI targets (**Figure 2d, 2e, and 2f**). These findings demonstrate the effectiveness of the PPI interface in conditioning the molecular generative model. For instance, MDM2/p53 and Bcl-2/Bax have notably different interface architectures and binding hot spots, which likely result in the generation of compounds with distinct QEPPI and Fsp3 profiles. These findings underscore the specificity of the conditioning framework, which adapts the molecular generation process to the target PPI interface, thus ensuring that the generated molecules are tailored to the unique features of each PPI target. Moreover, the drug-like properties of the generated compounds shifted relative to those in the training dataset, indicating that the GENiPPI framework does more than merely reproduce the distributions of known molecules; it generates novel compounds that maintain drug-likeness while exploring new regions of chemical space. This capacity to innovate within the bounds of known drug-likeness properties is a hallmark of successful generative models, as it enables the discovery of potentially more effective or optimized PPI modulators.

### Model performance

To assess the performance of the GENiPPI framework and compare it with other molecular generative models. We benchmarked our method using the MOSES platform[66], a leading benchmark platform of molecular generation. We trained all models on the full training dataset and randomly sampled 30,000 molecules. The models and hyperparameters provided by the MOSES platform were used, including the Adversarial Autoencoder (AAE)[67], character-level recurrent neural networks (CharRNN)[68], Variational Autoencoder (VAE)[69], LatentGAN[70] and ORGAN[71]. To validate the quality of the molecules generated by the conditioned model, we compared them to molecules generated by the GENiPPI framework and the GENiPPI-noninterface framework, which lacks the conditioned module. Our results showed that molecules generated by the conditioned GENiPPI framework outperformed those generated by other models in terms of novelty and diversity. This improvement can be attributed to the conditioning module, which directs the model to focus on specific chemical spaces associated with PPI interfaces. By conditioning molecular generation on relevant PPI features, the GENiPPI framework ensures that the generated molecules possess characteristics desirable for PPI inhibition. In contrast, models lacking this module, such as the GENiPPI-noninterface, fail to maintain this level of focus, resulting in lower performance across novelty and diversity metrics.

As shown in **Table 2**, the GENiPPI framework demonstrated advantages in uniqueness, novelty, and diversity over the GENiPPI-noninterface framework. Overall, the GENiPPI framework performed better in molecular generation. Compared to LatentGAN and ORGAN, GENiPPI offered superior results in terms of validity and diversity. While each molecular generative model has its strengths across various performance metrics, models tailored to specific tasks, such as those based on PPI structures, show advantages and inspirations from the GENiPPI framework.

**Table 2.**
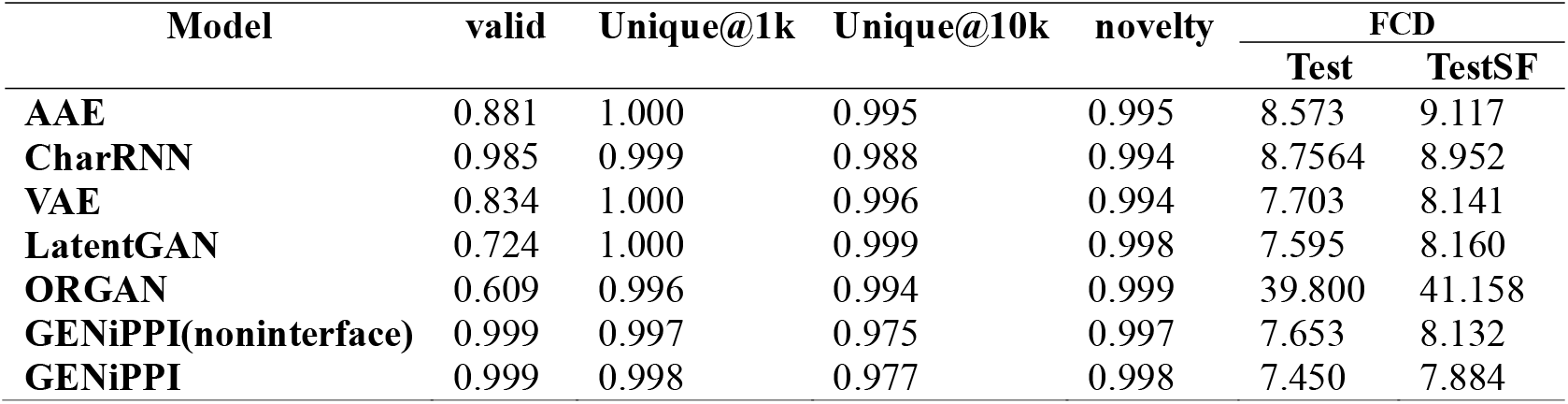
Valid, unique, novelty and FCD of sampling SMILES after training. We sampled 30,000 SMILES each time.

| Model | valid | Unique@1k | Unique@10k | novelty | FCD |  |
| --- | --- | --- | --- | --- | --- | --- |
|  |  |  |  |  | Test | TestSF |
| **AAE** | 0.881 | 1.000 | 0.995 | 0.995 | 8.573 | 9.117 |
| **CharRNN** | 0.985 | 0.999 | 0.988 | 0.994 | 8.7564 | 8.952 |
| **VAE** | 0.834 | 1.000 | 0.996 | 0.994 | 7.703 | 8.141 |
| **LatentGAN** | 0.724 | 1.000 | 0.999 | 0.998 | 7.595 | 8.160 |
| **ORGAN** | 0.609 | 0.996 | 0.994 | 0.999 | 39.800 | 41.158 |
| **GENiPPI(noninterface)** | 0.999 | 0.997 | 0.975 | 0.997 | 7.653 | 8.132 |
| **GENiPPI** | 0.999 | 0.998 | 0.977 | 0.998 | 7.450 | 7.884 |

To further understand the similarities and differences between the molecular distributions generated by the GENiPPI framework and other models, we compared the distribution of molecular properties in the Testset, iPPI-DB inhibitor[72], and the generated molecular datasets from AAE, CharRNN, VAE, LatentGAN, GENiPPI-noninterface and GENiPPI (**Supplementary Figs.5**). The generated compounds showed similar distributions of physicochemical properties to those in the training set. The QED values of the generated molecules were particularly close to those of the iPPI-DB inhibitors, indicating that the GENiPPI framework effectively learns and applies the desired molecular characteristics. Notably, while most iPPI-DB inhibitors have QED values lower than 0.5, the majority of generated molecules from the GENiPPI framework exhibited QEPPI values higher than 0.5. This suggests that the GENiPPI framework not only captures the drug-likeness of molecules but also their PPI-targeting potential, which is essential for PPI-related drug discovery tasks. This consistency in property distribution underscores the framework’s ability to model complex chemical spaces and generate biologically relevant, novel compounds.

### Chemical space exploration

To more comprehensively estimate the chemical space distribution of the model generated molecules in comparison to the active compounds from the training datasets, we evaluated the chemical drug-like space of the generated compounds by calculating t-distributed stochastic neighbourhood embedding (t-SNE) maps of MACCS fingerprints [73]. t-SNE is a widely-used dimensionality reduction method used for visualizing data points in two or three-dimensional space by mapping high-dimensional data to lower dimension[74, 75]. This method clusters similar compounds, allowing for a clear understanding of how generated compounds occupy chemical space in comparison to active, known compounds.

The distribution of both generated and active compounds in the chemical drug-like space was visualized using t-SNE visualization (**Figure 3a, 3b, and 3c**). Our findings reveal that the generated drug-like compounds not only share the chemical space with the active compounds, but are also homogeneously mixed in the two-dimensional space. This observation indicates that the GENiPPI framework successfully generates compounds that occupy the same drug-like space as known active modulators, reinforcing the model’s capacity to produce viable drug candidates. Moreover, under the 2D topological fingerprints, the generated compounds exhibit a similar chemical drug-like space to that of the active. This similarity suggests that the generative model is effective in capturing key topological features of molecules that are critical for drug-likeness. However, relying solely on two-dimensional representations may not be sufficient to fully assess drug-likeness, particularly for PPI modulators, which often require more complex three-dimensional features for effective binding. Incorporating three-dimensional features into the compounds contributes to the design of promising drug-like compounds[30, 76]. To address this, we conducted principal moments of inertia (PMI) shape analysis on the generated compounds and compared them with drug-like compounds from DrugBank and iPPI-DB(**Figure 3d**). This analysis revealed that many approved compounds are either rod or disk shaped, and the generated drug-like compounds display a similar three-dimensional shape distribution. Such shapes are often crucial for the spatial complementarity required for targeting PPI interfaces. Additionally, the PMI shape analysis suggests that the model is generating compounds with appropriate three-dimensional features that align with known drug-like shapes, further validating the robustness of the generation process. We also evaluated the plane of best fit (PBF) score of the generated drug-like compounds, a parameter that describes the extent to which molecular scaffolds deviate from planarity. The PBF distribution of the generated library ranges from 0 to 2 Å(**Figure 3e**), indicating that many generated drug-like compounds are derived from relatively planar molecular scaffolds.

Additionally, we assessed the ability of the model to generate PPI target-specific compounds using chemical space maps. To evaluate the overlap of drug-like chemical space, we utilized Tree MAP (TMAP) to create a 2D projection (**Figure 3f**)[77]. Each point represents a compound, colored by its target label, with dark and light colors denoting generated and active compounds, respectively. These results suggest that the GENiPPI model can generate compounds similar to the active compounds in the training set while introducing novel structures. Overall, the framework enriches and expands the chemical space of PPI-targeted drug-like compounds.

### Few-shot molecular generation

Due to the high associated with data collection, only a small amount of labeled biomedical data is typically available, presenting challenges for drug design and optimization, which often faces the problem of limited data[78]. This scarcity of labeled data can diminish the practical performance of most deep learning frameworks in drug design. Addressing the challenge of generating molecular designs with limited labeled data has become a significant focus in the few-shot generative community[79, 80]. Few-shot learning aims to train models using only a small number of examples while still enabling them to generalize effectively to novel tasks. This capability is critical for drug discovery, where only a few experimentally verified molecules are often available for new PPI targets. The GENiPPI model was applied to generate a virtual compound library targeting the interaction interface between heat shock protein 90 (Hsp90) and cell division cycle 37(Cdc37). By training the model on the PPI structure of Hsp90-Cdc37 (PDB ID: 1US7) and using data from seven disruptors, we sampled 500 valid compounds. The chemical space similarity between active disruptors and generated compounds for Hsp90/Cdc37 was visualized using t-SNE projection maps(**Figure 4a**), which revealed that the generated molecules were largely clustered around the active disruptors in chemical space. This result demonstrates the effectiveness of few-shot learning in navigating through the targeted chemical space and generating compounds that are structurally similar to known active disruptors despite limited training data.

In order to further assess the chemical relevance of the generated compounds, we performed pharmacophore-based matching using DCZ3112, a novel triazine derivative that disrupts Hsp90-Cdc37 interactions, as a reference molecule[81]. The top 5 generated molecules showed similar pharmacophore and shape features to DCZ3112(**Figure 4b**. The similarity of these features between DCZ3112 and the generated molecules indicates that the model was able to successfully learn the essential characteristics required for disrupting the Hsp90-Cdc37 interaction, even with limited input data.

To further validate the model’s performance, we examined the interaction patterns between the generated compounds and the PPI interface by performing molecular docking. Previous studies have identified key hot spot amino acid residues at the PPI interface (PDB ID: 1US7)[82, 83],as shown in **Figure 4c**. We performed molecular docking for prediction of the binding poses (**Figure 4d**) of DCZ3112 with the Hsp90-Cdc37 complex using the UCSF DOCK6.9 program[84]. The structure of the Hsp90-Cdc37 complex with DCZ3112 highlights the hydrogen bond interactions with amino acid residues: Arg32A, Glu33A, Ser36A, Ser115A, Gly118A, Gln119A, and Arg167B(**Figure 4d**), which are major contributors to protein-ligand interactions. Subsequently, molecular docking was also performed on the GENiPPI-generated compounds, alongside DCZ3112, and compounds with reasonable binding modes and higher binding affinity were selected for interaction pattern analysis. The GENiPPI-generated compounds not only achieved a better docking score than the active compounds but also reproduced the key interactions with the crucial residues of the PPI interface. This indicates that the model was able to accurately capture the binding preferences of the target PPI interface based on limited input data. The generated compounds formed additional halogen bonds, salt bridges and π-cation interactions, which were not observed in the reference disruptor. These additional interactions likely contributed to the improved binding affinity to the target PPI interface(**Figure 4e**). In conclusion, by analyzing the interaction patterns between the generated compounds and the PPI interface, GENiPPI successfully learned the implicit interaction rules between active compounds and the PPI interface. The ability to generate molecules that introduce novel interaction patterns, while retaining key interactions, demonstrates the model’s capability to innovate within the chemical space and design compounds with enhanced binding potential.

## Conclusion

In this work, we developed the GENiPPI framework, which combines PPI interface features with a conditional molecular generative model to generate novel modulators for PPI interfaces. Through extensive conditional evaluation experiments, we validated the ability of the GENiPPI framework to learn the implicit relationship between PPI interfaces and active molecules, demonstrating its capacity to generate chemically diverse and biologically relevant molecules. One of the key innovations of GENiPPI is its use of GATs to extract fine-grained, atomic-level interaction features from PPI interfaces. This allows the model to focus on critical interaction “hot spots” that are often difficult to target using conventional drug design methods. Additionally, by incorporating a conditional wGAN, the model is able to impose specific constraints on molecular generation, ensuring that the generated molecules are not only structurally novel but also align with the required PPI-targeting drug-likeness. Our comparative benchmarks and evaluation experiments across various settings demonstrate the practical potential of GENiPPI for rational PPIs drug design.

Despite these promising results, our framework has some limitations that can be addressed to improve its performance and applicability. First, the model has not been extensively tested across a large number of receptor-ligand pairs of PPIs, which may affect its generalization ability. This limitation arises from the relatively scarce data on drug-PPI target complexes compared to traditional drug-target dataset. The limited number of high-quality datasets for drug-PPI target complexes compared to traditional drug-target datasets poses a challenge. This scarcity of data remains a common issue in PPI-targeted drug discovery, largely due to the complexity of characterizing PPIs experimentally, and underscores the need for more comprehensive and curated datasets. Furthermore, the current framework does not incorporate the 3D structural information of ligand-receptor interactions in PPIs. Additionally, improvements can be made in representation learning, balancing training speed, and enhancing the diversity of generated molecules.

Several potential directions could further improve GENiPPI and its application to PPI-targeted drug discovery: (1) collecting and cleaning higher quality data pairs on receptor-ligand PPI complexes is essential. Improving the diversity and accuracy of the data used for model training can significantly enhance the model’s ability to generalize to new, unseen targets. (2) integrating molecular chemical language models and pre-trained models of protein-protein structural features to fine-tune receptor-ligand PPI datasets, thereby enhancing model generalization, novelty and diversity of generated compounds. (3) The current framework could be further enhanced by integrating fragment-based molecular generative models with 3D structural information. (4) Modifying the model architecture or combining it with deep reinforcement learning to optimize the generated compounds [85, 86]. By defining specific objectives such as maximizing binding affinity or improving pharmacokinetic properties, reinforcement learning agents could iteratively refine the generated

molecules to achieve more desirable drug-like characteristics.

In light of these potential improvements, our future work will focus on enhancing the GENiPPI framework by combining advanced representation learning methods and deep generative approaches.

In summary, the GENiPPI framework represents a significant advance in the field of PPI structure-based molecular generation. Its ability to integrate PPI interface features into a generative framework, combined with its performance in both few-shot learning and conditional evaluation experiments, highlights its potential as a powerful tool for rational PPI drug design.

## Methods

### Datasets

We first investigated PPI targets with sufficient compound bioactivity data for training and evaluation our model[87]. For this study, we selected 10 validated PPI drug targets that cover the binding interface (**Supplementary Tables S1**). These targets include E3 ubiquitin-protein ligase Mdm2, apoptosis regulator Bcl-2, BAZ2B, apoptosis regulator Bcl-xL, BRD4 bromodomain 1 BRD4-1, CREB-binding protein (CREBBP), ephrin type-A receptor 4 (EphA4), induced myeloid leukemia cell differentiation protein Mcl-1, and menin. Additionally, we randomly selected a subset of 250,000 compounds as additional inactive compounds from the ChEMBL dataset that was used as part of the training datasets[88]. A detailed data preprocessing can be found in **Supplementary Note A**.

### Model strategy and training

#### Graph attention networks of protein-protein interaction interface

In this section, the representation learning of protein-protein complex interfaces is inspired by previous work on protein docking model evaluation[60], which introduced a double-graph representation to capture the interface features and interactions of protein-protein complexes (**Supplementary Figs.1**). The extracted interface region is modeled as two graphs (*G*^1^ and *G*^2^), representing the interfacial information and the residues involved in the two interacting proteins. A graph *G* is defined as *G* = (*V, E*, and *A*), where *V* is the set of nodes, and *E* is the set of edges between them, and *A* is the adjacency matrix, which maps the association between the nodes, numerically representing the connectivity of the graph. If the graph *G* has *N* nodes, the dimension of the adjacency matrix A is ***N***\****N***, where **A**_***ij***_ **>0** if the *i*-th node is connected to the *j*-th node, and **A**_***ij***_ = 0 otherwise.

The graph ***G***^**1**^ encodes the atomic types of all residues in the interface region, and its adjacency matrix ***A***^**1**^ classifies the interatomic bonding types for all residues at the interface region. This representation only considers the covalent bonds between atoms of interface residues within each subunit as edges. The definition follows as:

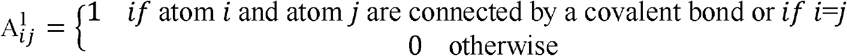

The graph ***G***^**2**^ represents both covalent bonds (including those captured ***G***^**1**^) and non-covalent residue interactions as edges. The adjacency matrix ***A***^**2**^ for ***G***^**2**^ accounts for both covalent bonds and non-covalent interactions between atoms that are within 10.0 Å of each other. The non-covalent atom pairs are defined as those whose distance is less than 10.0 Å. The definition follows as:

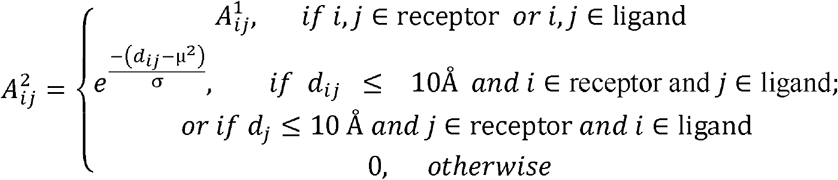

Here, ***d***_***ij***_ represents the distance between the *i*-th and the *j*-th atoms of all residues in the interaction region. *μ* and *σ* are learnable, with initial values of 0.0 and 1.0, respectively. The function 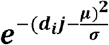 decays as the distance between atoms increases.

The graph representation provides a flexible and intuitive way to encode interactive information and adjacent(local) relationships. For the node features, we considered the physicochemical properties of the atoms, using the same features as in previous work[60, 89, 90]. The initial feature vector of each node, with a length of 23, was then embedded into 140 features using a one-layer fully connected (FC) network.

The constructed graphs are used as the inputs for the Graph Attention Networks (GATs). Each graph consists of adjacency matrices ***A***^**1**^, ***A***^**2**^, node matrices 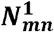, 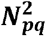, and the node features, 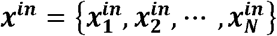 and ***x*** ∈ ℝ ^***F***^, where F is the dimensionality of the node features. For the input graph of ***x***^***in***^, the pure graph attention coefficients are defined as follows, representing the relative importance between the *i*-th and *j*-th nodes:

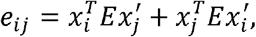

Here, 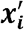 and 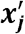 are the transformed feature representations, defined as 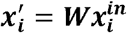 and 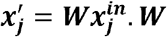, ***E*** ∈ ℝ ^**F ×F**^ are learnable matrices in the GATs. To satisfy the symmetrical property of the graph, ***e***_***ij***_ and ***e***_***ji***_ become identical by adding 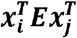 and 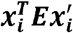. The attention coefficient will only be computed for *i* and *j* where ***A***_***ij***_ **> 0**.

The attention coefficients are also calculated for the elements in the adjacency matrix. For the element pairs (*i*,*j*), they are defined in the following form:

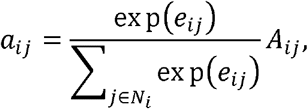

Here, ***a***_***ij***_ represents the normalized attention coefficient between the *i*-th and *j*-th node pairs, while ***e***_***ij***_ is the computed symmetric graph attention coefficient. ***N***_***i***_ denotes the set of neighbors for the *i*-th node, which includes the interacting node *j* with ***A***_***ij***_ **> 0**. The goal is to define attention by simultaneously considering both the physical structure ***A***_***ij***_ and the normalized attention coefficient ***e***_***ij***_ of the interactions.

Based on the attention mechanism, the new features of each node are updated by considering its neighboring nodes. This update is a linear combination of the neighboring node features and the final attention coefficient ***a***_***ij***_ :

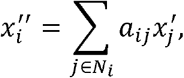

Using the previously described GATs mechanism, we applied four layers of GATs to process the node embedding information from the neighboring nodes and output the updated node embedding. For the two adjacency matrices ***A***^**1**^ and ***A***^**2**^, we use a shared GAT. the initial input to the network consists of the atomic feature. With two matrices ***A***^**1**^ and ***A***^**2**^, we compute ***x***_**1**_ **= *GAT* (*x***^***in***^, ***A***^**1**^ **)and *x***_**2**_ **= *GAT* (*x***^***in***^,***A***^**2**^).

To focus exclusively on the intermolecular interactions at the interface of the input protein-protein complex, we obtain the final node embedding by subtracting the embeddings of the two graphs. By subtracting the updated embedding ***x***_**1**_ from ***x***_**2**_, we can capture aggregated information about intermolecular interactions from the other nodes at the protein-protein complex interface. The output node feature is therefore defined as:

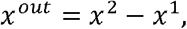

Afterward, the updated *x*^*out*^ becomes *x*^*in*^ and iteratively passes through the subsequent three GAT layers to further increase the information. After all four GAT layers updated the node embeddings, the embedding of all nodes in the graph are summed to represent the overall intermolecular interaction of the protein-protein complex:

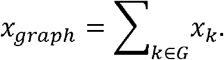

Finally, fully connected (FC) layers were applied to the ***x***_***graph***_ to obtain a [4,4,4] features vector representing the protein-protein interface.

#### Molecular representation

For each SMILES string, a 3D conformer is generated using RDKit[91] and optimized with the default settings of the MMFF94 force field. The molecular structure is then extracted into a 35Å grid centered at the geometric center of the molecule using the HTMD package[92]. The atoms are discretized into a 1 Å cubic grid, and eight channels are used to compute voxelized information. Finally, the electronic density for the 9th channel is calculated using the original molecule method in Multiwfn(**Supplementary Figs.2**) [93].

#### Conditional Wasserstein generative adversarial networks

The generator takes a conditional vector and a noise vector sampled from a Gaussian distribution as inputs. The PPI interface features([1,4,4,4], vector shape) are concatenated with a noise vector of size [9, 4, 4, 4] and input into a 4-layer transposed convolutional neural network (CNN) with 256, 512, 1024, and 1024 filters, respectively. The first three layers downsample the array size using strided convolution (stride=2). For all convolutions, a kernel size of 4 is used, and the Leaky ReLU is applied as the activation function after each convolution. BatchNorm3d is applied between the convolution and activation operations to normalize the values across each channel for each sample.

The discriminator consists of a 4-layer sequential convolutional neural network (CNN) with 256, 512, 1024, and 1024 filters, respectively. The first three layers downsample the array size using strided convolution (strided=2). As with the generator,, a kernel size of 4?is used for all convolutions, and Leaky ReLU (α=0.2) is applied as the activation. InstanceNorm3d is applied between the convolution and activation steps to normalize the values across each channel for each sample.

The physical and spatial features of the compounds are derived from the molecular representation learning module, while the PPI interface features are obtained from the GATs module of the protein complex interface. These features are used to estimate the matching probability between molecules and the PPI interface features(**Supplementary Figs.3**).

#### Molecular captioning network

In this section, we describe the process of decoding the generated molecular representation into SMILES strings. Our approach is inspired by shape-based molecular generation[94, 95], which utilize a combination of convolutional neural networks (CNNs) and Long Short-Term Memory (LSTM) networks to generate SMILES strings. Briefly, the molecular captioning network consists of a 3D CNN and a recurrent LSTM networks. The molecular representation generated by the generator is first fed into the 3D CNN, and the output is then passed into the LSTM to decode the SMILES strings (**Supplementary Figs.4**).

#### Model training

The conditional generative adversarial network is trained using Wasserstein loss. The loss functions for the generator (G_(0(z,c))_), and discriminator (D_0_ (x)) are as follows:

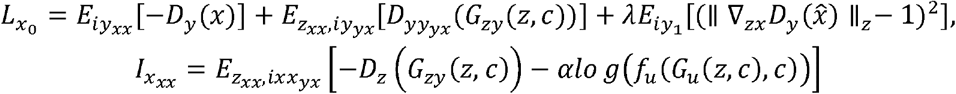

Here, x and *c* represent molecular representations and PPI interface features, respectively, sampled from the true data distribution p_real_. The variable *z* is a random noise vector sampled from a Gaussian distribution (*p*_*z*_), and *f*_0_ is a function that evaluates the probability that a PPI interface feature corresponds to a molecular representation. The terms λ and α are regularization parameters, both empirically set to 10. The λ term controls the effect of the gradient penalty on discriminator loss, while the α term controls the influence of *f*_0_ on the generator’s loss.

The model was trained for 50,000 iterations with a batch size of 8 (65 steps per iteration). The discriminator was updated after each step, while the generator was updated every 30 steps. The network was trained using the RMSprop optimizer, with a learning rate of 1 × 10^-4^ for both the generator and discriminator. During training, we monitored the similarity between real and generated molecular representations using Fréchet distances. The weights of the conditional networks were pre-trained using binary cross-entropy loss and were frozen during GAN training. Training was performed on a single NVIDIA A40 GPU, and all neural networks were built and trained using Pytorch 1.7.1[96] and Tensorflow 2.5[97].

#### Molecular generation

After training, the embedding information of the protein-protein complex interface is used to guide the model in generating novel molecules from the latent space. The maximum sampling strategy was applied in the LSTM, where the next token in the SMILES string is generated by selecting the one with the highest prediction probability[94].

### Evaluation settings

#### Conditional evaluation metrics

In this study, the primary objective was to evaluate the effectiveness of the proposed framework for protein-protein interaction (PPI) interface-based conditional molecular generation. We sampled the same number of valid molecules for three PPI targets. For the generated compounds and comparison sets, we calculated the QED and Fsp3 values using RDKit, and the QEPPI values using the QEPPI package (https://github.com/ohuelab/QEPPI). The density distribution of these drug-likeness metrics was then plotted to compare the differences.

#### MOSES evaluation metrics

To evaluate the performance of our proposed conditional molecule generation framework, we used the evaluation metrics of validity, uniqueness, novelty and diversity provided by the MOSES platform, which are defined as follows: Validity: Molecules defined as valid in the generated molecules.

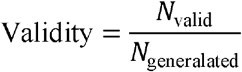

Uniqueness: The proportion of unique molecules found among the generated valid molecules.

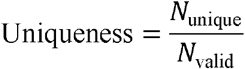

Novelty: The generated molecules are not to be covered in the training set.

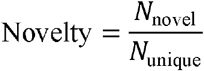

FCD(Fréchet ChemNet Distance):To detect whether the generated molecules are diverse and whether they have chemical and biological properties that are similar with the real molecules[98].

#### Molecular shape

To evaluate the shape space of molecules, we used two widely adopted molecular descriptors to represent the three dimensions of molecular structure: principal moment of inertia (PMI)[99] and the best-fit plane (PBF)[100]. The PMI descriptor classifies the geometric shape of molecules based on the degree to which they are rod-shaped (linear shape, such as acetylene), disk-shaped (planar shape, such as benzene), or sphere (spherical shape, such as adamantane). The normalized PMI ratios (NPRs) are plotted on a two-dimensional triangle to compare the shape space covered by different sets of molecules, allowing for the evaluation and visualization of the diversity of molecular shape with a given set. The PBF descriptor is a three-dimensional measure that represents the deviation of a molecule from a plane. It is defined as the mean distance of each heavy atom from the best-fit plane passing through all heavy atoms.

#### Tree MAP

To explore and visualize the chemical space through unsupervised visualization of high-dimensional data, we calculated MinHash fingerprint vectors for both active and generated compounds[101]. We then used tmap and faerun to construct two-dimensional projections using Tree MAP (TMAP)[102].

#### Protocol for few-shot generation

Targeting the Hsp90-Cdc37 PPI interface is recognized as an important strategy for cancer therapy. The crystal structure of the Hsp90-Cdc37 protein complex (PDB ID: 1US7) is available for molecular docking[103]. In addition, known Hsp90-Cdc37 PPI disruptors were collected for training in few-shot generative tasks. These disruptors include DCZ3112, Celastrol, FW-04-804, Sulforaphane, Withaferin A, Platycodin D, Kongensin A [104].

OpenPharmacophore(https://github.com/uibcdf/OpenPharmacophore) was utilized to create pharmacophore models and perform virtual screening. The protein structures were processed using UCSF Chimera[105], the program DOCK6.9 was used for semiflexible docking. Figures were generated using PyMOL[106]. A detailed docking protocol is provided in **Supplementary Note B**.

## Abbreviations

PPIs: protein-protein interactions
cWGAN: conditional wasserstein generative adversarial network
CNNs: convolutional neural networks
GATs: graph attention networks
LSTM: long short-term memory
QED: quantitative estimation of drug-likeness
QEPPI: quantitative estimate of protein-protein interaction targeting drug-likeness;
Fsp3: fraction of sp3 carbon atoms
t-SNE: t-distributed stochastic neighbor embedding.

## Data availability

The original data downloaded from ChEMBL and processed datasets are available on Zenodo (https://zenodo.org/records/13968592). All the source code and datasets are available at Github (https://github.com/AspirinCode/GENiPPI).

## Acknowledgements

This research was supported by the Yonsei University graduate school “Integrative Biotechnology”.

## Author information

### Authors and Affiliations

Department of Integrative Biotechnology, Yonsei University, Incheon 21983, Republic of Korea

Jianmin Wang, Jiashun Mao & Kyoung Tai No

School of Informatics, Yunnan Normal University, Kunming, China

Chunyan Li

College of Computer Science and Electronic Engineering, Hunan University, Changsha, Hunan, 410082, China

Hongxin Xiang & Xiangxiang Zeng

School of Computer Science and Technology, China University of Petroleum, Qingdao, 266580, Shandong, China

Xun Wang, Shuang Wang & Tao Song

Department of Computer Science, University of Tsukuba, Tsukuba, 3058577, Japan Zixu Wang & Yangyang Chen

College of chemistry and chemical engineering, Lanzhou University, Lanzhou 730000, China

Yuquan Li

High Performance Computer Research Center, University of Chinese Academy of Sciences, Beijing, 100190, China

Xun Wang

## Contributions

J. Wang collected data, developed the model, analyzed the data, and wrote the manuscript; J. Mao and C. Li developed the model and analyzed the data; H. Xiang, X. Wang, S. Wang, Z. Wang, Y. Chen, Y. Li helped to refine the research through constructive discussions and revised the manuscript; K. No, T. Song, X. Zeng supported and supervised the project, interpreted the results, and wrote revisions to the manuscript.

## Corresponding author

Correspondence to Xiangxiang Zeng; Tao Song; Kyoung Tai.

## Ethics declarations

Competing interests

The authors declare no competing interests.

## Supplementary Information

Method details: Supplementary A, Dataset and preprocessing; Supplementary B, Docking protocol; Supplementary C, Model evaluation; Supplementary Tables S1, .

Overview of the 10 PPI interface datasets used for training; Supplementary Figures S1, GATs module for representation learning of the protein complex interface; Supplementary Figures S2, CNNs module for molecular representation learning; Supplementary Figures S3, cWGAN module for conditional molecular generation; Supplementary Figures S4, molecular captioning network module for SMILES strings decoding; Supplementary Figures S5, Distribution of molecular properties.

